# Prelimbic Cortex Activity Predicts Anxiety-Like Behavior in the Elevated Plus Maze

**DOI:** 10.1101/2024.12.26.630448

**Authors:** Marina A. Smoak, Karla J. Galvan, Daniel E. Calvo, Rosalie E. Powers, Travis M. Moschak

## Abstract

The medial prefrontal cortex (mPFC) plays a critical role in emotional regulation, and its dysregulation is linked to anxiety disorders. In particular, the prelimbic cortex (PrL) of the mPFC is thought to modulate anxiety-related behaviors, though its precise role remains debated. Here, we used endoscopic in vivo calcium imaging to assess PrL neuronal activity in male and female Sprague-Dawley rats performing in the Elevated Plus Maze (EPM), a widely used task to measure anxiety-like behavior. We found that animals that spent less time in the open arms exhibited higher PrL activity in the open arms, suggesting that heightened PrL activity may reflect greater anxiety or increased avoidance behavior. These results suggest that the PrL may play a role in regulating the emotional response to anxiety-provoking situations, potentially influencing the tolerance for exposure to threatening environments.

## Introduction

The prefrontal cortex (PFC) integrates corticolimbic inputs to evaluate emotional significance and guide adaptive behavior (McLaughlin *et al*., 2014). Variations in PFC activity are linked to reported anxiety levels (Simpson *et al*., 2001), and dysregulation of PFC activity is associated with mental disorders such as generalized anxiety disorder (GAD) (Cha *et al*., 2014), post-traumatic stress disorder (PTSD) (Shin *et al*., 2005), and social anxiety (Stein *et al*., 2002). Given these findings, it is essential to isolate and evaluate the activity of the PFC to clarify how normal emotional responses become dysregulated in anxiety-inducing conditions.

The human PFC is a large region consisting of several subregions. Functional magnetic resonance imaging (fMRI) studies have demonstrated that participants diagnosed with generalized anxiety disorder (GAD) showed sustained activation of the anterior cingulate cortex [Brodmann area (BA) 32; ACC] (Paulesu *et al*., 2010). In a recent fMRI study that evaluated individuals in an approach-avoidance task, high-anxiety participants relied on areas 24/25 (ACC) for emotional-action control, whereas their non-anxious peers did not (Bramson *et al*., 2023). In rodents, the prelimbic cortex (PrL) of the mPFC is considered homologous to the ACC (areas 24/32) of non-human primates (Heilbronner *et al*., 2016). Both inactivation (Stern *et al*., 2010; Green *et al*., 2020) and activation (Wang *et al*., 2015) of the PrL during elevated plus maze (EPM) testing has been shown to increase open arm exploration, suggesting an ambiguous and context-dependent role for the PrL in regulating anxiety-related behaviors in rodent models. To shed light on what might be driving these disparate findings following PrL manipulation, we investigated PrL neural activity in the EPM task.

Specifically, this study assessed anxiety-like behaviors in male and female Sprague-Dawley rats using the EPM, a well-established paradigm for measuring anxiety phenotypes (Hogg, 1996). During behavior, we used endoscopic in vivo Ca^2+^ imaging to measure neuronal responses in the PrL. We found that increased PrL activity in the open arms was associated with less time spent in open arms.

## Methods

### Subjects and Surgery

Female (n=5) and male (n=10) adult Sprague Dawley rats aged 8–10 weeks and weighing 200–300 grams were singly housed in a temperature-controlled room (21 C1 C) under a 12-hour light/dark cycle (lights on at 8 PM) with food and water ad libitum. Upon arrival to the facility, animals were given 7 days to acclimate, followed with at least 7 days of handling by experimenters. All experiments were conducted during the dark phase. All procedures were conducted following the National Institutes of Health Guidelines for the Care and Use of Laboratory Animals and in accordance with the University of Texas at El Paso Institutional Animal Care and Use Committee.

Rats underwent stereotaxic surgery for intracerebral injection of the viral vector encoding the genetically coded calcium sensor GCaMP6s (pAAV.Syn.GCaMP6s.WPRE.SV40/100843-AAV5) and lens implantation in the prelimbic area (AP: +2.7; ML: +/-0.6 DV: -3.6 for virus, -3.4 for lens). Infusions were counterbalanced for hemisphere (left or right). To secure the implantation, four stainless steel skull screws (Small Parts) were fastened to the skull, two rostrally and two caudally. After the lens was implanted and the skull thoroughly dried, fresh fast curing orthodontic acrylic resin powder and liquid (Lang Dental) were applied. Once completely hardened, a small protective boundary was created around the lens using dental cement.

During surgery, animals were anesthetized with a cocktail of 100 mg/kg ketamine and 10 mg/kg xylazine and were given a one-week recovery period. Post-operative care included daily meloxicam and 10% enrofloxacin diluted in sterile saline. After one week of recovery, skin margin care was performed to ensure proper wound healing and to prevent infection. If fluorescence was detected in the brain after ∼4–6 weeks, subjects then underwent installation of Miniscope baseplates and implantation of an intrajugular catheter with the port externalized at the back. Post-operative care ensued with treatments of cefazolin, meloxicam, and daily flushing with 2% heparinized saline. Following recovery, animals ran in behavioral tasks involving operant responding to sucrose pellets that were unrelated to the current study, followed by the elevated plus maze assessment mentioned below.

### Elevated Plus Maze and Behavior

A standard elevated plus maze constructed with black acrylic and equipped with infrared sensors at the entrance of each runway (Med Associates) was utilized to measure anxiety-like behaviors. The maze was placed in a silent and dimly lit room, elevated 199.94 cm from the ground, and consisted of two open arms (58.8 x 10.16 cm) and two closed arms with no roof (50.8 x 10.16 x 40.29 cm) at right angles to each other, and an open square in the center (10.16 x 10.16 cm). During this task, rats were placed in the center square with their face pointing toward a closed arm of the elevated plus maze. The experimenter left the room, and subjects were left alone for 10 minutes. During this time, subjects were allowed to move freely within the apparatus. Infrared photobeam sensors placed along the walls of the maze determined the position of animals while in the elevated plus maze.

### Endoscopic In Vivo Ca^2+^ Imaging

Imaging was recorded using software and hardware from the Miniscope system (UCLA Miniscope, Los Angeles, CA) with time-locked task events. Specifically, the Miniscope camera was attached to the implanted baseplate on the animal’s head and connected via the Miniscope data acquisition (DAQ) QT Software (https://github.com/Aharoni-Lab/Miniscope-DAQ-QT-Software). The DAQ system also included a transistor-transistor logic (TTL) connection to Med-PC (v.5), which controlled all behavioral systems and would trigger recordings captured using a high-powered Dell laptop. The program recorded neuronal activity for the entire 10 minutes of the task. The animal was removed from the maze and placed back in its home cage for 10 minutes of recording. This additional recording ensured enough neural activity was obtained for subsequent data analysis.

### Data Analysis

Videos of calcium activity were obtained from Miniscope software, and time spent in the open arms versus the closed arms of the elevated plus maze was obtained through Med-PC. As described by Sullivan & Sdrulla (2022), raw imaging files within the same field of view were concatenated into single, shorter videos containing timestamps using the Miniscope DAQ QT Software. Raw imaging files were assessed for quality before analysis and downsampled before being analyzed in CaImAn software (Pnevmatikakis et al., 2016; Giovannucci et al., 2019). Using CaImAn’s built-in rigid motion correction algorithm, downsampled videos were median filtered and motion corrected in chronological order. Putative neurons were identified, restricted by a convolutional neural network classifier, and separated utilizing CaImAn’s nonnegative factorization method. This method simultaneously accounts for the spatial and temporal components of calcium signals. The activity of each putative neuron was estimated using the deconvolution algorithm within CaImAn. This algorithm contains parameters set to match the kinetics of GCaMP6s (Chen *et al*., 2013). All putative neurons were manually inspected and artifactual or noisy neurons were excluded from further analysis. Once neural activity was detected in CaImAn, Matlab was used to align behavioral events that were captured by MedPC to neural activity. Extracted estimated activity trains were segmented according to whether they occurred while the animal was in the open arms, closed arms, or junction. We then calculated the relative neural activity per zone (open arms, closed arms, junction) over the total activity for the session as a measure of “Neural preference” (e.g., open arm activity / total activity). Two male animals spent less than 20s in the open arms and were thus excluded from the analysis as there was too little activity to accurately analyze. Statistical analysis was performed using IBM SPSS Statistics 29.0. Statistical significance level was set to α=0.05. Unpaired *t*-tests were used to compare sex, percent time spent in open arms and open arm entries. Median splits were performed on behavioral data such as number of open arm entries and percent time spent in open arms. Pearson correlations were used to compare neural activity to number of open arm entries and the percent time spent in open arms.

### Histology

Upon completion of the experiment, animals were euthanized, the brain was carefully removed, left overnight in 4% paraformaldehyde, and transferred to 30% sucrose the following day. Subsequently, the brains were set in optimal cutting temperature embedding medium (O.C.T., Fisher) and frozen on a tissue mount/specimen chuck in a cryostat (Leica CM3050 S Cryostat), sectioned coronally at 30 μm, and mounted on slides to verify lens placement. Verification was visualized via widefield fluorescence using a Zeiss LSM 900 with Airyscan 2 confocal microscope and placement was mapped onto an atlas (Paxinos and Watson, 2007).

## Results

### Prl neural activity predicts time spent in open arms

Animals were classified using a median split into groups with low or high percent time spent in the open arms and low or high number of open arms entries. Subjects in the Low group could be considered to have higher anxiety-like behavior and subjects in the High group could be considered to have lower anxiety-like behavior. Median splits were also considered for percent time spent in closed arms and percent time spent in the junction of the elevated plus maze. Unpaired *t*-tests verified that groups were significantly different when evaluating for differences in percent time spent in open arms (**Fig. 1A**, t(10) = 6.376, *p* < 0.001) and number of open arm entries (**Fig. 1B**, t(10) = 5.966, *p* = 0.001). Groups were then further evaluated for differences in prelimbic activity. Subjects who spent less time in the open arms had a significantly higher proportion of PrL activity in the open arms when compared to subjects who spent more time in the open arms (**Fig. 1C**, t(10) = 3.062, *p* = 0.01). No differences in PrL activity were seen when comparing the number of open arm entries (**Fig. 1D**, t(10) = 0.3061, *p* = 0.76). Furthermore, subjects who spent more time in the closed arms had a significantly higher proportion of PrL activity in the closed arms when compared to subjects who spent less time in the open arms (*p* = 0.007) but not the percent time spent in the junction (*p* = 0.58). When comparing all subjects, Pearson’s correlations revealed a relationship between PrL activity and percent time in open arms (*r* = –0.56, *p* = 0.04; **Fig. 1E**), where higher PrL activity was associated with more anxious behavior. We saw no relationship between PrL activity and the number of open arm entries (*r* = –0.30, *p* =0.3160; **Fig. 1F**). Furthermore, no significant relationship was seen between percent time spent in closed arms (*r* = 0.51, *p* = 0.06) or percent time spent in the junction (*r* = –0.37, *p* = 0.21).

**Figure 1.**
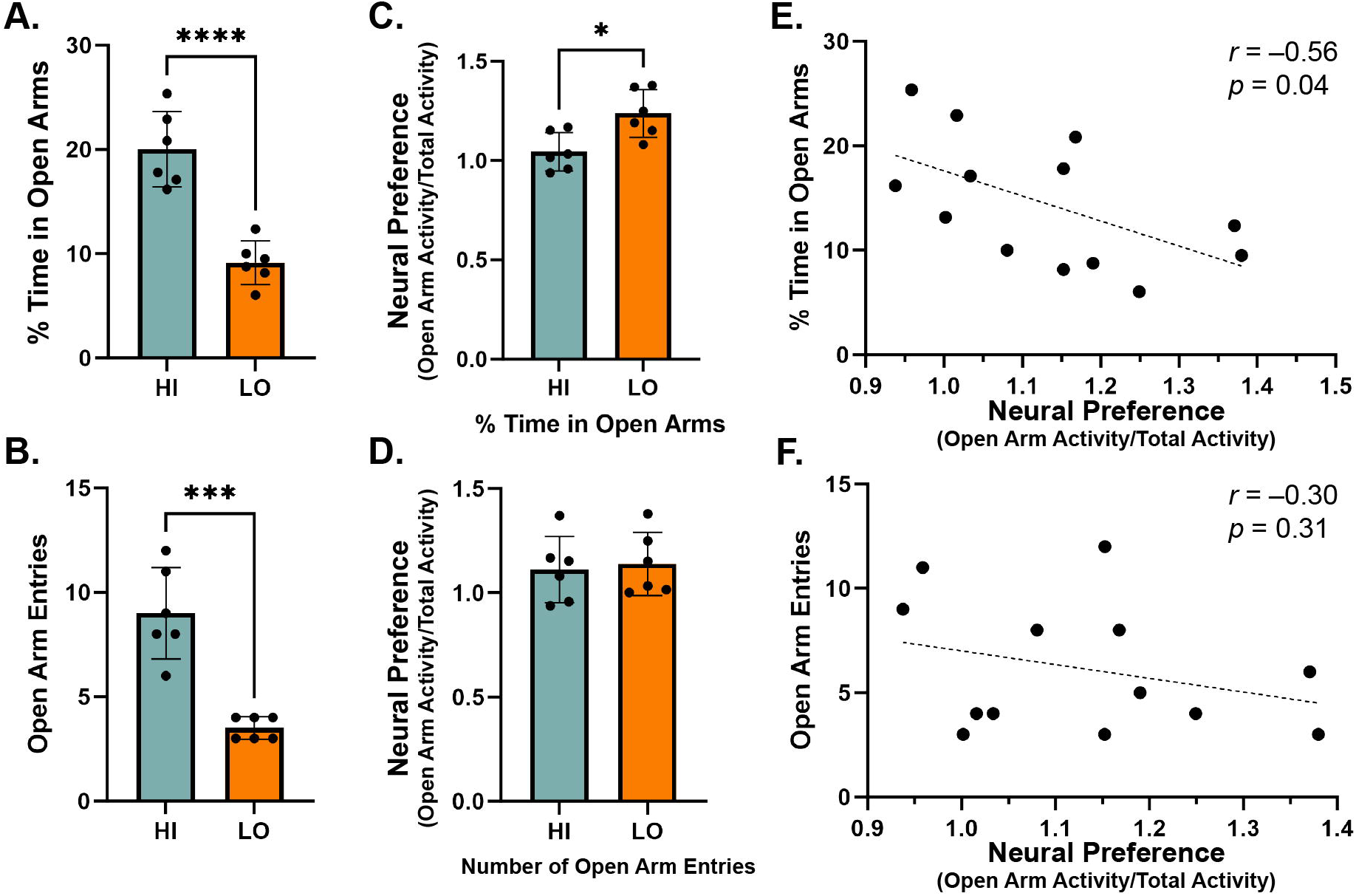
Behavioral and neural predictors of anxiety-like behaviors. Median splits were used to classify two groups of subjects where lower anxiety-like behavior was determined by a greater percent time spent in open arms or more open arm entries (HI) and increased anxiety-like behavior was determined by a lesser percent time spent in open arms or fewer open arm entries (LO). Unpaired t-tests confirmed that the groups differed significantly in terms of the percentage of time spent in the open arms (A, p < 0.05) and the number of open arm entries (B, p < 0.05). When groups were evaluated for differences in neural activity, subjects who spent less time in the open arms exhibited a significantly higher percentage of PrL activity compared to those who spent more time in the open arms (C, p < 0.05), but not number of open arm entries (D, p = 0.76). Pearson’s correlations across all subjects revealed a negative relationship between PrL activity and the percentage of time spent in the open arms (E), indicating that higher PrL activity was associated with greater anxiety. No relationship was found between PrL activity and the number of open arm entries (F). * p ≤0.05; *** p ≤0.001; **** p ≤0.0001

### Histology

Data were only included if cells and lens placement were verified to be in the prelimbic cortex. The placements and locations for all subjects included in this study can be seen in **Figure 2**.

**Figure 2.**
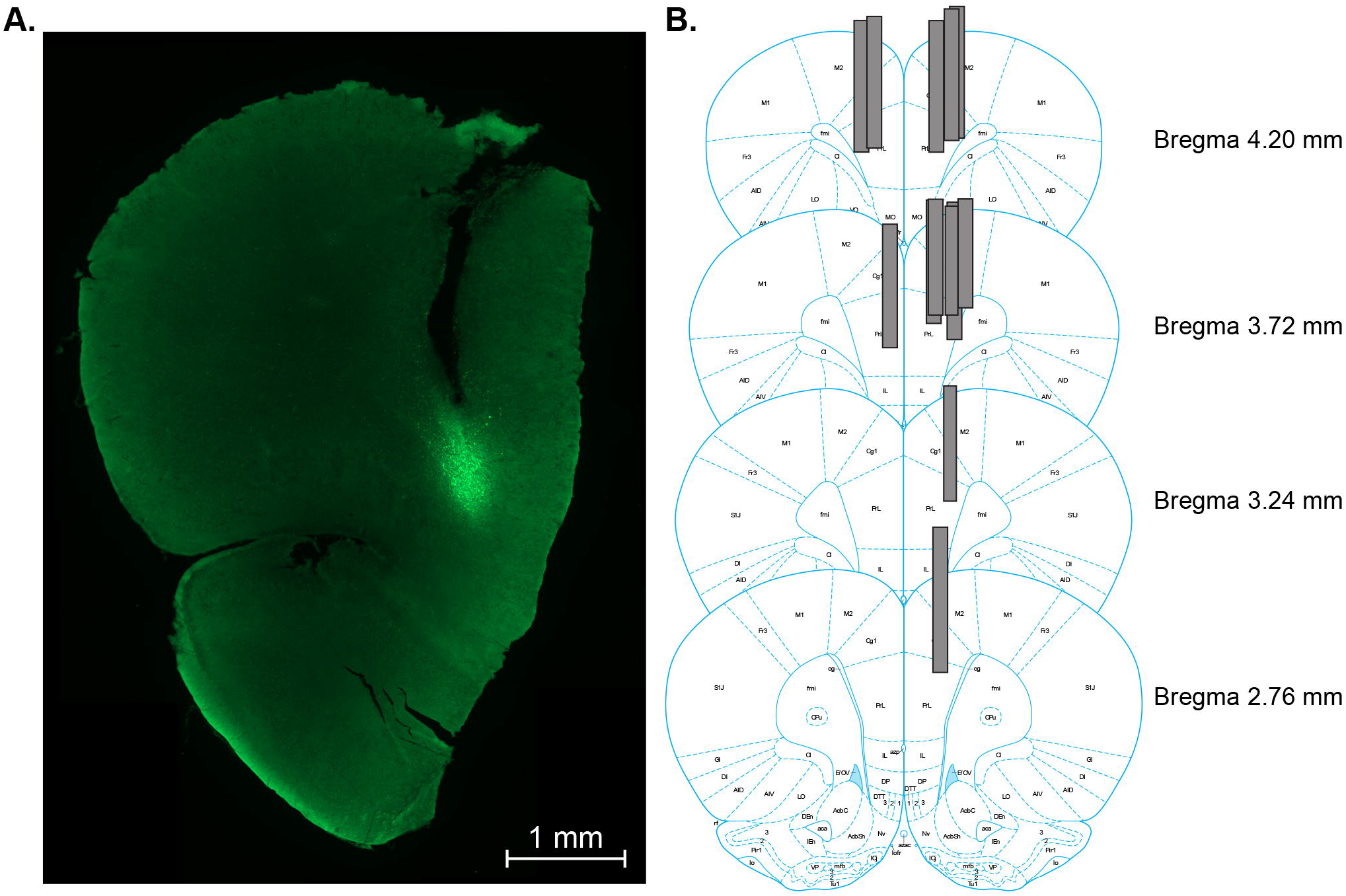
Rat coronal maps displaying the locations of lens implantation for the animals used in this study. Using coronal sections of fluorescent images (A), lens placement was neuroanatomically verified and subsequently mapped onto templates of a rat brain atlas (B, Paxinos & Watson, 2006).

## Discussion

Overall, our findings demonstrate that high PrL activity in the open arms predicts increased anxiety-like behavior in the EPM. However, no correlation was found when comparing PrL activity with the number of open arm entries. These results suggest that PrL activity may more closely associate with the duration of time spent in exploration of the open arms rather than the decision to enter the open arms. Since the amount of time spent in the open arms of the EPM may reflect the subject’s level of anxiety or willingness to remain in an exposed, potentially threatening environment, PrL neural activity could be linked to the animal’s tolerance of/coping with anxiety (Aliczki *et al*., 2016), or greater exploration despite fear (Casanova *et al*., 2024; Marin-Blasco *et al*., 2024) in this paradigm.

Our findings reinforce many previous preclinical outcomes. For example, temporary PrL inactivation during EPM testing increased open arm exploration (Stern *et al*., 2010; Green *et al*., 2020). Additionally, lesions of the entire mPFC (including the PrL) yielded a higher percentage of open arm entries and time spent in open arms (Lacroix *et al*., 2000). Furthermore, effects of dorsal and ventral mPFC infusions of the benzodiazepine midazolam produced anxiolytic effects as determined by greater open arm entries and percentage of time in open arms (Shah *et al*., 2004). Finally, optogenetic suppression of the PrL-BLA pathway in chronic pain mice increased time spent in open arms (Gao *et al*., 2023). However, these effects may depend on neural subtype and receptor. Notably, activation of mPFC β-adrenoceptors in excitatory neurons was anxiogenic (Lei *et al*., 2022), and intra PrL injection of a selective dopamine D4 receptor antagonist increased time spent in the open arms (Vergara et al., 2017), while activation of either Ca^2+^/calmodulin-dependent protein kinase α (CamKIIα)-positive excitatory neurons in the PrL (Pati *et al*., 2018) or *Drd1* expressing neurons in the mPFC produced anxiolytic responses (Hare *et al*., 2019). Furthermore, some studies have found null or even opposing effects of PrL manipulation on anxiety-like behavior. Stimulation of the PrL had no effect on EPM behavior in one study (Shimizu *et al*., 2018), while optogenetically activating the contralateral PrL to an inflamed hind paw produced anxiolytic effects (Wang *et al*., 2015). Furthermore, reversible inactivation of bilateral ventral portions of the mPFC (including the PrL) enhanced anxiety during EPM testing as determined by decreased number open arm entries (Lisboa *et al*., 2010). Nonetheless, our findings support the bulk of studies manipulating the PrL that suggest that heightened activity in the PrL is associated with heightened anxiety-like behavior (Lacroix *et al*., 2000; Stern *et al*., 2010; Green *et al*., 2020; Gao *et al*., 2023).

Few studies have assessed PrL neural activity during the EPM. In one, recording of multiunit activity via wireless telemetry revealed no change in the firing rate of PrL neurons during open arm entries, indicating that PrL neurons may not play a role in overcoming anxiety-like behaviors (Shimizu *et al*., 2018). While our results did not replicate these findings, other studies have provided evidence that there are dynamic changes in PrL activity between safe and aversive locations in the EPM. Lu et al., (2018) reported strengthened theta oscillations in PrL functional connectivity and increased information transfer efficiency while in the closed arms of the EPM. This further supports that anxiety-induced changes in neural activity occur in the PrL and enhanced connectivity may contribute to the inhibition of exploratory behaviors, which the PrL is known to encode (Ahmadlou *et al*., 2021; Brockway *et al*., 2023).

In addition to rodent work, clinical studies suggest that individuals with anxiety show increased ACC activation, especially when tasked with emotional regulation or threat anticipation (Amir *et al*., 2005; Simmons *et a*l., 2008; Paulesu *et al*., 2010; Maier *et al*., 2012; Fullana *et al*., 2016; Bramson *et al*., 2023; Buehler *et al*., 2024), although activation patterns vary depending on the specific type of anxiety disorder or task (Burkhouse *et al*., 2018). These clinical findings, which highlight increased ACC activation in anxiety contexts, align with our preclinical observations and those of others, suggesting that similar neural circuits, particularly within the mPFC, may play a central role in the regulation of anxiety across species.

## Conclusion

We found that subjects with higher anxiety-like behavior exhibited increased PrL activity in the open arms of the EPM. These results validate previous findings and strengthen the reliability of the prelimbic cortex’s role in anxiety across different methodologies and experimental conditions. Future studies could focus on the dynamics of this relationship by incorporating real-time tracking of activity in PrL projections implicated in EPM. One pathway of interest is the PrL’s projection to the basolateral amygdala, as optogenetic suppression of PrL neurons projecting to the basolateral amygdala increased the time spent in open arms (Gao *et al*., 2023). The ventral hippocampus is an alternative target of interest that projects to the PrL, since in the retrieval of fear extinction memory in rodents, the primary input to active neurons in the PrL came from the ventral hippocampus (Szadzinska *et al*., 2021). Finally, it would be valuable to examine how individual differences, such as genetic factors within the PrL (Chen *et al*., 2017) or it’s inputs (Hallock *et al*., 2020), or prior stress exposure (Corcoran *et al*., 2007; Quiñones-Laracuente *et al*., 2021; Smiley *et al*., 2021), modulate PrL activity or connectivity and anxiety responses, which may help tailor more personalized approaches to anxiety treatment.

## Acknowledgements

This work was supported by National Institute on Drug Abuse (NIDA) grant DA045764 and National Institute of Health (NIH) grant U54MD007592-28 awarded to TMM and the Dr. Keelung Hong Graduate Research Fellowship awarded to MAS. The authors would also like to thank Peter A. Fogel for the valuable discussions.

## Author Contributions

**Marina A. Smoak**: Data Curation, Formal Analysis, Visualization, Writing-original draft, Writing-review and editing; **Karla J. Galvan**: Investigation, Project Administration, Visualization, Writing-original draft; **Daniel E. Calvo**: Investigation, Writing-original draft; **Rosalie E. Powers**: Investigation, Visualization, Writing-original draft; **Travis M. Moschak**: Conceptualization, Formal analysis, Funding acquisition, Supervision, Writing-review and editing.

## Data Accessibility

The data presented in this report are available upon reasonable request from the corresponding authors.

